# Single-nucleosome imaging reveals principles of transient multiscale chromatin unfolding triggered by histone ADP-ribosylation at DNA lesions

**DOI:** 10.1101/2024.08.28.610034

**Authors:** Fabiola García Fernández, Catherine Chapuis, Junwoo Park, Eva Pinto, Victor Imburchia, Edoardo José Longarini, Angela Taddei, Nataliya Sokolovska, Ivan Matić, Sébastien Huet, Judith Miné-Hattab

## Abstract

Timely access to DNA lesions is crucial for genome integrity. This process requires profound remodeling of densely packed chromatin to establish a repair-competent architecture. However, limited resolution has made it impossible to fully understand these remodeling events. Here, combining microirradiation with live-cell multiscale imaging, we report that DNA damage-induced changes in genome packing rely on the conformational behaviour of the chromatin fiber. Immediately after damage, a transient increase in nucleosome mobility switches chromatin from a densely-packed state to a looser conformation, making it accessible to repair. While histone poly-ADP-ribosylation is required to trigger this switch, mono-ADP-ribosylation is sufficient to maintain the open-chromatin state. The removal of these histone marks by the ARH3 hydrolase then leads to chromatin recondensation. Together, our multiscale study of chromatin dynamics establishes a global model: distinct waves of histone ADP-ribosylation control nucleosome mobility, triggering a transient breathing of chromatin, crucial for initiating the DNA damage response.

## INTRODUCTION

The high level of DNA packing displayed by chromatin in the cell nucleus represents a major challenge for DNA-transaction processes including the repair of genetic alterations. The DNA damage response (DDR) is characterized by multiple chromatin remodeling processes. Among them, histones tails undergo post-translational modifications (PTMs) and remodeling^1^, facilitating efficient and faithful genomic restoration. One of the earliest remodeling events is the rapid and transient relaxation of the chromatin architecture occurring within seconds after DNA damage induction to facilitate access to the lesions^2–6^. This rapid unfolding is triggered by ADP-ribosylation (ADPr), a modification known to contribute to several repair pathways, such as DNA strand breaks resolution^7^. Upon recruitment to DNA lesions, the polymerase PARP1 adds ADP-ribose marks on nearby proteins, primarily PARP1 itself and histones ^8^. While this signaling pathway has been usually considered as mainly composed of poly-ADP-ribose (PAR) chains, recent findings evidenced a distinct, more enduring mono-ADP-ribose (MAR) signal, potentially displaying distinct functions^9,10^. The homeostasis of these two components of the ADPr pathway is controlled by HPF1, a PARP1 cofactor regulating its catalytic activity^11–13^, as well as hydrolases preferentially targeting PAR or MAR marks^14–17^. While PARG is the most active PAR hydrolase, ARH3 is a specific serine MAR eraser^15^. PARP1 has been identified as a central regulator of the chromatin architecture for several decades^7^. *In vitro*, PARP1 binding to nucleosomes was reported to promote the compaction of isolated chromatin fibers^18^ while PARP1 catalytic activity was rather involved in chromatin fiber unfolding^19^. More recently, live-cell experiments have shown that PARP1-dependent histone ADPr regulates chromatin compaction state in the vicinity of DNA breaks^4,20,21^. Nevertheless, it remains unknown how this modulation of chromatin compaction relates to conformational changes at the level of the chromatin fiber. As for any polymer, there is an intimate relationship between the chromatin architecture and its dynamics, although the exact characteristics of this relationship remain only partially understood^22^. Multiple studies have reported an increase in chromatin dynamics upon DNA damage, indicative of major changes in the underlying chromatin architecture^23–32^. Such increase in chromatin dynamics is now considered as an integral part of the DDR, for example favoring homology search during homologous recombination (HR)^30,33,34^. However, this compelling model originates mainly from studies in yeast, leaving the actual picture in mammalian cells more ambiguous^35^. Depending on the kind of DNA damage, the distance from the lesions as well as the time after damage induction, various impacts on the local dynamics of the chromatin fiber have been reported in mammalian systems^23,28,36–38^. More importantly, previous studies have traditionally focused on single spatial scales, preventing the establishment of a comprehensive model for changes in chromatin structure in the DNA damage response.

In the present work, we combine single-molecule imaging and micro-irradiation in human cells to dissect the remodeling events undergone at different scales of the chromatin structure during early steps of the DDR. This multiscale approach enabled us to discover that, within the first seconds after DNA damage, a temporary increase in nucleosome mobility alters chromatin from a densely packed state to a looser conformation, making it accessible to the repair machinery. Moreover, building on the recent surge of new insights into ADPr signaling, we demonstrate that histone ADPr is a master regulator of these remodeling events, with differential roles played by the PAR and MAR signals. Our findings provide a solution to the decades-long puzzle of reconciling the transient nature of poly-ADPr with the enduring effect of PARP1 on chromatin by assigning an open chromatin maintenance function to histone mono-ADPr.

## RESULTS

### Multi-scale chromatin remodeling occurs immediately after DNA damage

In order to get a comprehensive view of chromatin behavior immediately after DNA damage, we assessed chromatin dynamics at three folding scales: the global compaction state, the chromatin fiber and the nucleosome. To measure changes affecting the compaction state, we irradiated the nucleus of Hoechst-presensitized human U2OS cells expressing H2B fused to the photo-activatable dyes PAGFP using a continuous 405 nm laser (Figure 1A). Such irradiation simultaneously induces DNA lesions and highlight the damaged area. We monitored the thickness of the photoconverted line to assess changes in the level of chromatin compaction. In agreement with our previous findings^4^, we observed a rapid chromatin relaxation at DNA damage sites peaking 1 minute after damage. Noteworthy, this rapid unfolding is not associated with significant nucleosome disassembly^4^, implying that it mainly relies on conformational changes undergone by the chromatin fiber. Following the rapid relaxation phase, chromatin remained in an open state for a few minutes and then slowly recondensed to ultimately reach a compaction level that is beyond the pre-damage state (Figure 1A), in line with previous findings ^39^.

**Figure 1.**
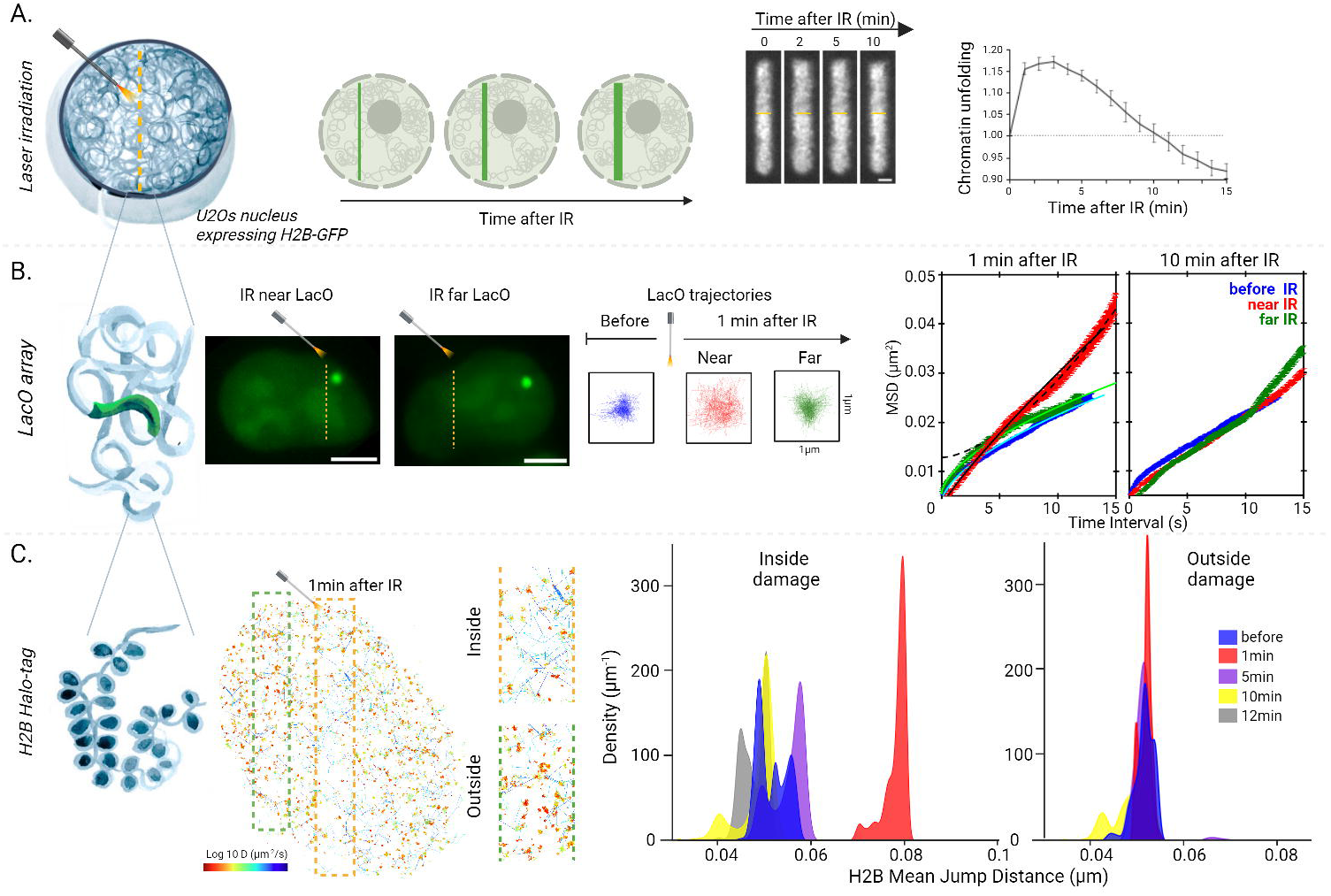
Chromatin undergoes rapid multiscale remodeling immediately at DNA lesions. On the left of each panel is shown a schematic representation of the chromatin folding scale that is monitored. (A) Nuclear scale. Sketch and representative confocal image sequence of a subregion of the nucleus of U2OS cells expressing H2B-PAGFP and presensitized with Hoechst after irradiation with a continuous 405 nm laser to simultaneously trigger DNA damage and photo-labeling of the irradiated region. Scale bar: 2 μm. The average thickness of the photo-activated line is plotted as a function of time after irradiation and normalized to time zero to estimate the changes in the overall chromatin compaction state (n=16). (B) Chromatin fiber scale. Representative images of U2OS nucleus harboring a fluorescently tagged *lacO* array and irradiated with a pulsed 355 nm laser nearby or away from the array. Scale bars: 8 µm. Representative locus trajectories and mean squared displacement (MSD) curves (n=10) before damage (blue) and 1 minute and 10 minutes after irradiation near (red) or far from the locus (green). Scale bar: 1 µm. Fitting of sub-diffusive and directed motion regimes is represented by solid and doted curves, respectively. (C) Nucleosome scale. Trajectories of individual histones in the nucleus of a U2OS cell expressing H2B-Halo bound to PA-JF549 Halo ligand. H2B motions were monitored 1 min after irradiation at 355 nm. A magnified view of the tracks inside and outside the irradiated area is shown on the right. The trajectories are color-coded according to their diffusion coefficient using the look-up table shown below. Mean jump distance histograms for the immobile population of H2B tracks inside (left) and outside (right) the irradiated region, before and at different times after micro-irradiation. Number of cells analyzed (N). Inside damage: N_bef_=52, N_1min_=35, N_5min_=20, N_10min_=26, N_12min_=28; Outside damage: N_bef_=20, N_1min_=7, N_5min_=11, N_10min_=11. Mean jump distance between each condition versus before damage are significantly different inside the irradiated region (p < 0.001, calculated from Yuen-Welch Test) but not outside (p > 0.05, calculated from Yuen-Welch Test).

Next, we analyzed how these rapid changes in the compaction state correlated with a modulation of the conformational behavior of the chromatin fiber. First, we assessed the dynamics of the fiber in cells harboring a *lacO*-array, inserted at a single genomic location in a euchromatic region of chromosome 1, visualized with GFP fused to LacI^40^. We monitored the dynamics of the tagged locus before, 1 minute and 10 minutes after DNA damage induced by irradiation with a pulsed 355 nm laser, nearby the locus or away from it (Figure 1B). The motion of the locus was quantified by computing the mean square displacement (MSD) curves from 10 individual trajectories (Figure 1B). In agreement with previous reports^41–47^, this analysis revealed a subdiffusive behavior prior to DNA damage (MSD(t) ∼0.004 t ^0.59^, R2 = 0.994) consistent with the Rouse model previously used to describe chromatin motion^48–50^. One minute following irradiation nearby the *lacO* array (≃1µm), we observed an increase in chromatin dynamics as shown by the higher amplitude of the MSD curve (Figure 1B, red curve). The fitting of the MSD revealed a complex diffusion behavior. While, at short time scales (t < 5 min), the motion remained subdiffusive although with a higher anomalous exponent (MSD ∼ 0.003 t ^0.9^, R2 = 0.997), at longer time scales, the locus rather exhibited a directed motion (MSD ∼ 0.0002 t ^1.9^, R2 = 0.998). Such directed motion at long timescale is consistent with chromatin decondensation that tends to push chromatin away from the irradiated area, as previously reported^4^. Ten minutes after damage, chromatin recovers its initial dynamics in correlation with its recompaction. In contrast, when irradiation was performed away from the fluorescent locus (≃8µm), the MSD followed anomalous diffusion (MSD ∼ 0.006 t ^0.5^, R2 = 0.994) similar to locus dynamics prior to DNA damage(Figure 1B, green curve). Altogether, this analysis reveals a striking increase in the dynamics of the chromatin fiber in the vicinity of the DNA damage region.

To increase the resolution of our analysis one step further, we monitored the impact of DNA damage induction on the dynamics of individual histones. Using H2B fused to HaloTag (H2B-Halo) and labeled with the photoactivable Janelia Fluor PA-JF549 HaloTag ligand ^51^, we combined laser micro-irradiation and single molecule imaging to follow the 2-dimensional trajectories of individual nucleosomes (Figure 1C, left). While almost no traces could be recovered in unlabeled control cells, in the presence of this construct, we obtained thousands of tracks per nuclei expressing H2B-Halo, with a mean track length of about 17 frames (Figure S1A-C, supplementary movie 1). In line with previous reports^52–56^, the tracks showed very limited motion of H2B proteins in the absence of damage, in contrast to freely diffusive HaloTag fused to a nuclear localization signal (Figure S1D). Using a convolutional neural network (CNN) deep-learning algorithm (Figure S1E, F), H2B trajectories were classified in 3 populations, similar to previous reports (Figure S1G and ^53,56–58^). The slow population, which gathers the vast majority of the tracks (83±5 %), most likely corresponds to H2B proteins stably incorporated into the nucleosomes as they display an effective diffusion coefficient similar to that of the chromatin fiber assessed with the *lacO* array (D_H2B_ =0.0012 μm^2^/s and D_lacO_ = 0.001 μm^2^/s). The mobile fraction (12±5 % of the tracks) shows a diffusion coefficient (D_H2B_ = 0.154 μm^2^/s) that is two-order of magnitude larger than the immobile population. Together with a hybrid population switching from immobile and mobile phases (5±2 % of the tracks), these fast trajectories are probably associated with the small fraction of histones that are not stably associated with the chromatin fiber and therefore rapidly diffuse within the nucleoplasm.

Then, we applied this analysis pipeline to study the behavior of individual H2B proteins in and out the area of DNA damage (Figure 1C). Of note, we controlled that histone mobility was not affected by the successive imaging sequences used to monitor H2B trajectories at different timepoints after DNA damage (Figure S1H). We found that the proportions of the different populations of tracks were only mildly impacted by damage induction (Figure S1G), in line with our previous observations showing no major nucleosome disassembly at these early steps of the DDR^4^. We focused our attention to the population of slow H2B proteins likely incorporated into the nucleosomes and assessed their mobility by measuring the mean distance covered in 10 ms for each H2B trajectory, a generic metric that did not require to assume a specific diffusion model. A rapid surge in mobility restricted to the irradiated region was observed, culminating in a ∼60% increase in nucleosome motion at 1 min post-damage (Figure 1C). This dramatic increase was only transient, with a rapid recovery as early as 5 min after irradiation, the nucleosomes becoming even less mobile than prior to damage at later timepoints. Comparing these data with the changes in the overall chromatin compaction state (Figure 1A) shows that the acute increase in nucleosome dynamics correlates with the rapid relaxation process. In contrast, chromatin remains in this decompacted state for several minutes despite the rapid drop in nucleosome dynamics. Therefore, while the increase in nucleosome mobility seems to underlie chromatin unpacking, it does not appear necessary for the maintenance of the open state.

### PARP1-mediated ADPr signaling is the central trigger of multiscale chromatin remodeling at DNA lesions

In line with the rapid recruitment of PARP1 at sites of laser irradiation (Figure S2A), we previously showed that ADPr signaling controls the early modulation of chromatin compaction state at DNA lesions^4^. Here, we investigated whether these changes in chromatin packing could be linked to ADPr-dependent remodeling events at the level of the chromatin fiber. We monitored nucleosome dynamics in the presence of the clinically-relevant PARP inhibitor (PARPi) Talazoparib that impedes the catalytic activity of PARP1 and leads to its prolonged retention at DNA lesions (Figure S2A), in line with previous observations^59^. While PARPi treatment slightly impacts the dynamics of the *lacO* array and the nucleosome in the absence of damage (Figure S2B), it led to decreased mobility in both assays after laser irradiation (Figure 2B, S2B). Therefore, PARPi treatment did not only suppress the enhanced motions of the chromatin fiber observed in untreated cells, but even reduced these motions. We also assessed chromatin dynamics in cells knocked out (KO) for PARP1, the main driver of ADPr signaling in the context of the DDR^60^. The loss of PARP1 suppressed the increased nucleosomes mobility observed in wild-type (WT) cells at sites of damage, but did not lead to reduced dynamics, regardless the presence of PARPi (Figure 2C-D). Therefore, while the recruitment of inhibited PARP1 restrains nucleosome motions at sites of damage, PARP1-dependent ADPr increases these motions. These findings nicely correlate with those regarding the global chromatin packing state, which showed that inactive PARP1 enhances chromatin compaction at DNA lesions while ADPr promotes unfolding^4^.

**Figure 2.**
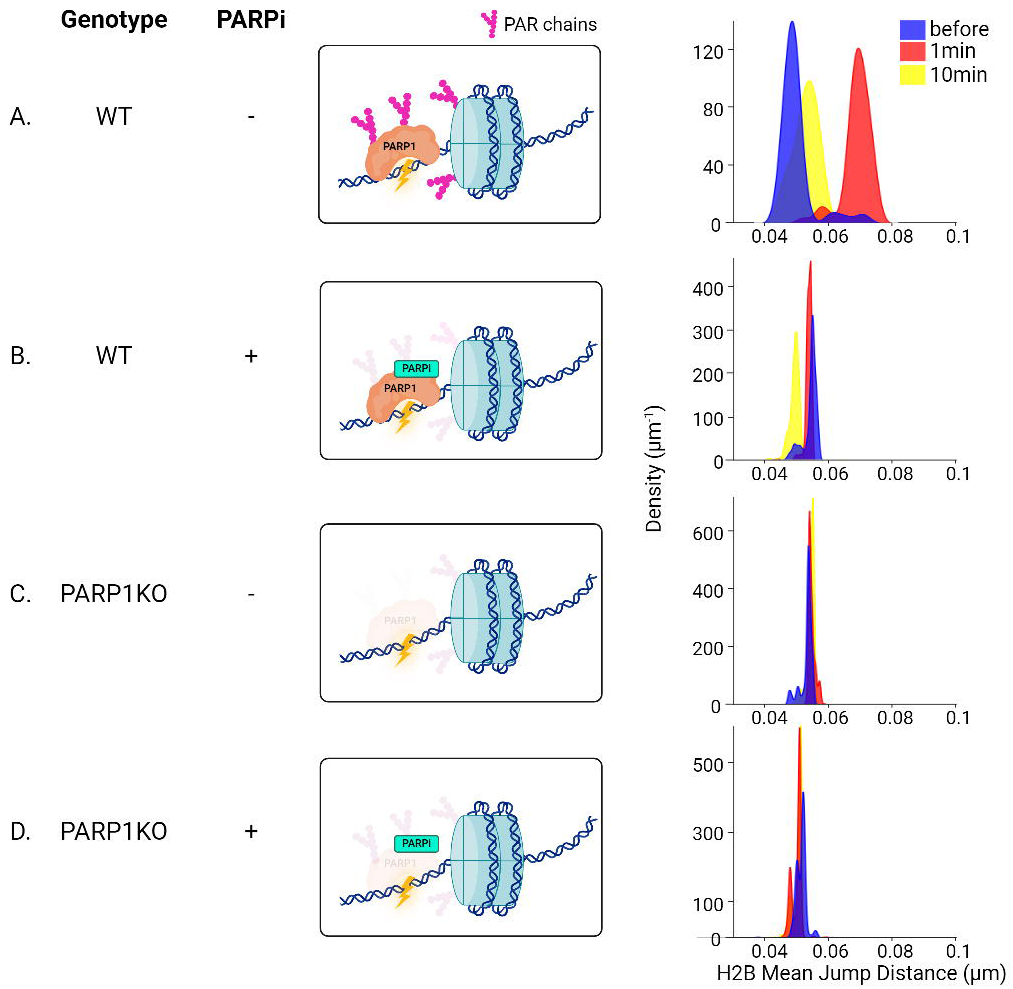
ADP-ribosylation by PARP1 triggers enhanced nucleosome dynamics after micro-irradiation. On the left of each panel, a sketch shows the status of the ADP-ribose signal depending on the U2OS genotype and PARPi treatment (30 µM Talazoparib). On the right is shown the mean jump distance histograms for the immobile population of H2B tracks inside the irradiated region, before and at different times after micro-irradiation at 355 nm in WT (A, B) and in PARP1KO (C, D). (WT N_bef_=12, n_1min_=4, N_10min_=3; WT + PARPi N_bef_=44, N_1min_=31, N_10min_=6; PARP1 KO N_bef_=13, N_1min_=12, N_10min_=8; PARP1 KO + PARPi N_bef_=10, N_1min_=9, N_10min_=10).

### Spontaneous increase in ADPr signaling is sufficient to increase chromatin fiber dynamics

Besides ADPr, the cell activates an intricate network of signaling pathways at sites of DNA damage^61^. Therefore, it is difficult to assign the changes in chromatin dynamics we observed at the lesions to a direct effect of ADPr rather than a potential crosstalk of this signaling with other DDR-related pathways. To assess the specific impact of ADPr on chromatin dynamics, we took advantage of cells lacking the hydrolase ARH3 that show spontaneous activation of ADPr signaling, in particular upon treatment with an inhibitor against the poly-ADP-ribose-glycohydrolase (PARGi) (Figure 3A). Importantly, this was not associated with enhanced γH2AX signaling, a classical responder of DNA breaks, showing that the strong ADPr signal observed in these cells is not the consequence of a global activation of the DDR^16^. Therefore, comparing WT and ARH3 KO cells treated or not with PARGi allows for assessing the specific impact of ADPr signaling on chromatin folding independently of the DDR context. We found that nucleosome dynamics was higher in ARH3 KO compared to WT cells and could be further enhanced by PARGi treatment (Figure 3B, C). These data indicate a clear correlation between the level of activation of the ADPr pathway and the dynamics of the chromatin fiber, independently of the presence of DNA lesions. Therefore, enhanced ADPr appears sufficient to promote the local mobility of the nucleosomes along the chromatin fiber.

**Figure 3.**
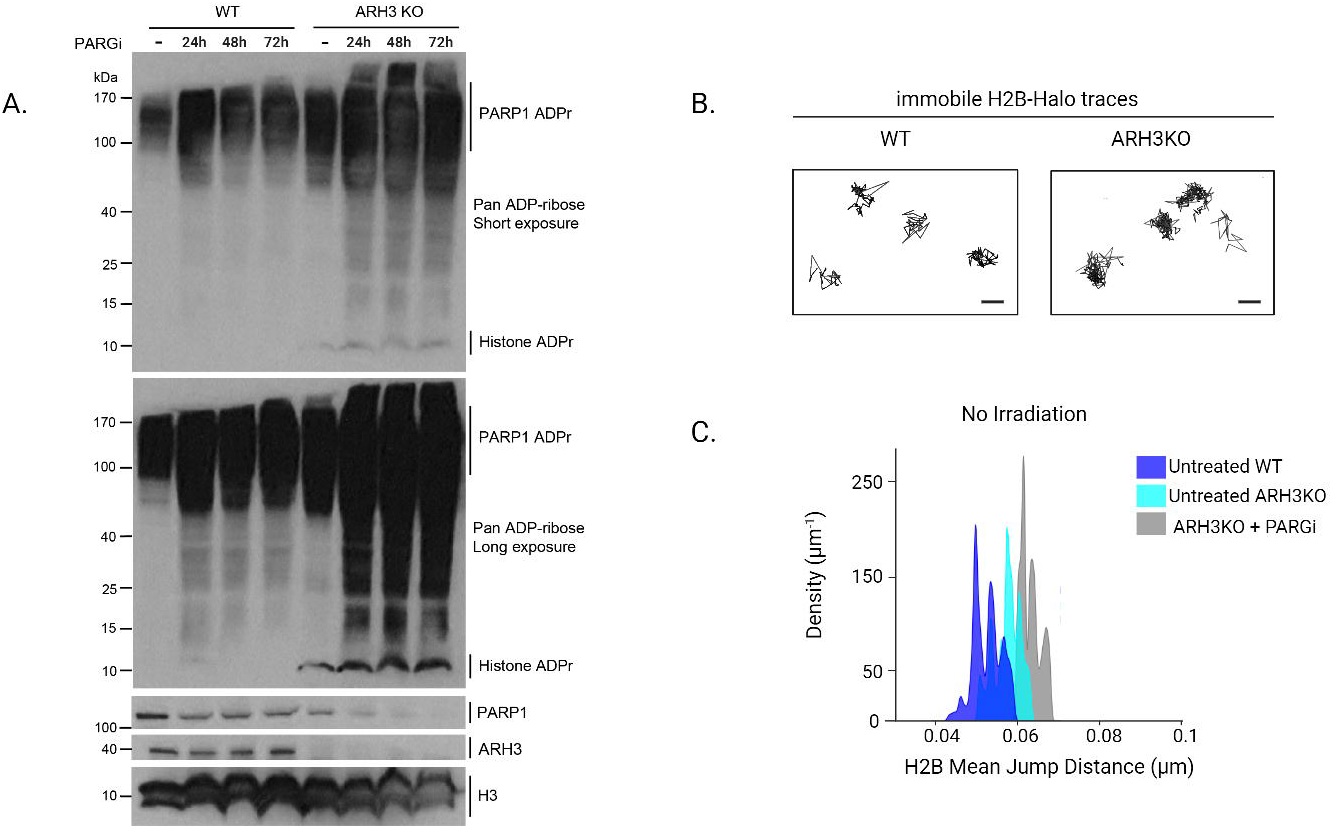
Spontaneous ADPr signal upon loss of ARH3 is sufficient to increase nucleosome dynamics. (A) Western blot displaying APDr signal, stained with a pan-ADPr antibody as well as PARP1 and AHR3 signals in WT or ARH3 KO left untreated or after 24 to 72 hs of PARGi treatment (25 µM PDD00017273). H3 is used as loading control. (B) Representative examples of the immobile population of H2B trajectories inside the nucleus of undamaged WT and ARH3 KO cells. (C) Mean jump distance histograms for the immobile population of H2B tracks in undamaged WT and ARH3 KO cells, treated or not with 25 µM PDD00017273 PARGi for 24 hs. (WT N=75, untreated ARH3 KO N=55, ARH3 KO + PARGi N=22). Mean jump distance between each condition *versus* WT condition are significantly different (p < 0.001, calculated from Yuen-Welch Test).

### Histone ADPr is needed to establish a dynamic open chromatin state at DNA lesions

Upon DNA damage, ADPr signal is found mainly on PARP1 and the different histones^8,17,62^. To disentangle the relative contributions of PARP1 automodification and histone ADPr on chromatin dynamics, we studied the impact of two PARP1 mutants. In the PARP1-3SA, the three main Ser residues targeted by ADPr (S499, S507, S519) are switched to Ala, leading to a strong decrease in automodification while not affecting histone ADPr^63,64^. Instead, the PARP1-LW/AA mutant (L1013A/W1014A) is unable to ADP-ribosylate histones due to impaired interaction with HPF1^21,65^. While the expression of wild type PARP1 or PARP1-3SA in PARP1 KO cells both rescued the transient increase in nucleosome mobility at sites of DNA lesions, this was not the case for PARP1-LW/AA (Figure 4). These data demonstrate that it is the ADPr of histone and not PARP1, that triggers the increase in chromatin fiber mobility at sites of DNA breaks. Together with the fact that histone ADP-r was also shown to control chromatin decondensation at the lesions^21^, our findings draw a model in which the addition ADP-r marks along the chromatin fiber increases its mobility, which itself promotes global unfolding.

**Figure 4.**
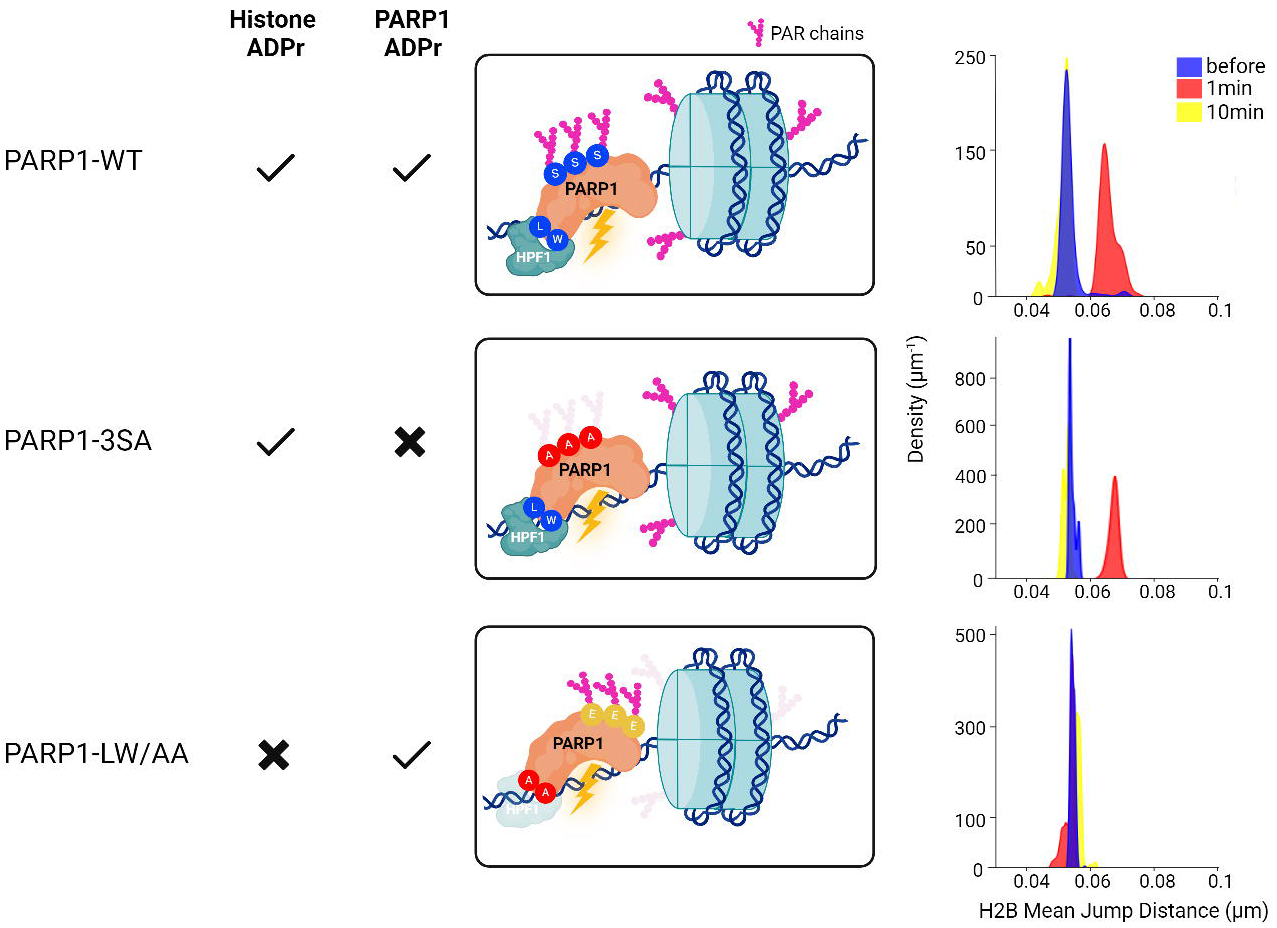
The transient increase in nucleosome dynamics at sites of damage is controlled by histone ADP-ribosylation. On the left of each panel, a sketch shows the characteristics of PARP1 automodification and histone ADP-ribosylation depending on the PARP1 construct expressed in U2OS PARP1 KO cells. On the right is shown the mean jump distance histograms for the immobile population of H2B tracks inside the irradiated region, before and at different times after micro-irradiation at 355 nm. (PARP1-WT N_bef_=13, N_1min_=6, N_10min_=4; PARP1-3SA N_bef_=29, N_1min_=9, N_10min_=13; PARP1-LW/AA N_bef_=14, N_1min_=12, N_10min_=8).

### The erasure of mono-ADP-ribose is needed for chromatin recondensation

Consecutively to its initial rapid unfolding, chromatin remained in an open state for a few minutes. This was followed by a slow recondensation phase which led to a compaction state that is higher than the pre-damage one (Figure 1). While chromatin opening was shown to be important for facilitating access to DNA lesions^21^, the recondensation was also proposed to trigger the recruitment of some members of the repair machinery, potentially in relation to transcription shut-down at sites of DNA lesions^39,66,67^. Given the key role played by ADPr signaling during the chromatin unfolding step, we wondered whether this pathway could also regulate the recondensation process.

First, we monitored the kinetics displayed by the ADPr signal to compare them to those of chromatin relaxation (Figure 1). In agreement with our recent findings^9^, we observed that ADPr signal at DNA lesions could be decomposed in an early acute PAR peak and a more progressive and sustained MAR wave (Figure 5A). The timeframe of these two components suggests that, while the transient PAR surge may trigger chromatin unfolding, the maintenance of the open state might be rather controlled by the more persistent MAR signal. To test this hypothesis, we analyzed the impacts of the loss of ARH3. Indeed, this hydrolase, while not regulating the PAR signal, controls the progressive removal of the MAR marks at DNA lesions (Figure 5A). Importantly, the loss of ARH3 also triggered a possible imbalance in the double-strand break repair pathways as shown by the increased accumulation of the NHEJ-related protein 53BP1 in ARH3 KO cells while the HR-related protein BRCA1 accumulation remained unchanged (Figure S3). Together with increased sensitivity to genotoxic stress observed in ARH3 KO cells^68^, these data suggest that the timely removal of MAR signaling by ARH3 contributes to efficient DNA repair. Regarding chromatin remodeling at sites of damage, we found that, while not affecting unfolding, the loss of ARH3 strongly impaired the recondensation process (Figure 5B). AHR3 KO cells did not reach the over-condensed state observed in WT cells and were even unable to recover to the pre-damage chromatin compaction level. At the single nucleosome scale, ARH3 KO displayed persistent increased nucleosome mobility up to 10 min post-irradiation in contrast to the recovery observed in WT cells (Figure 5C). These different findings reveal that the erasure of the MAR signal by ARH3 is crucial for the restoration of the chromatin structure following its early unfolding upon damage induction. This provides the first functional explanation for the recently revealed temporal bimodality of PARP1 signaling^9^ and deepens our understanding of the key role played PARP1 in the control of chromatin structure at sites of DNA damage.

**Figure 5.**
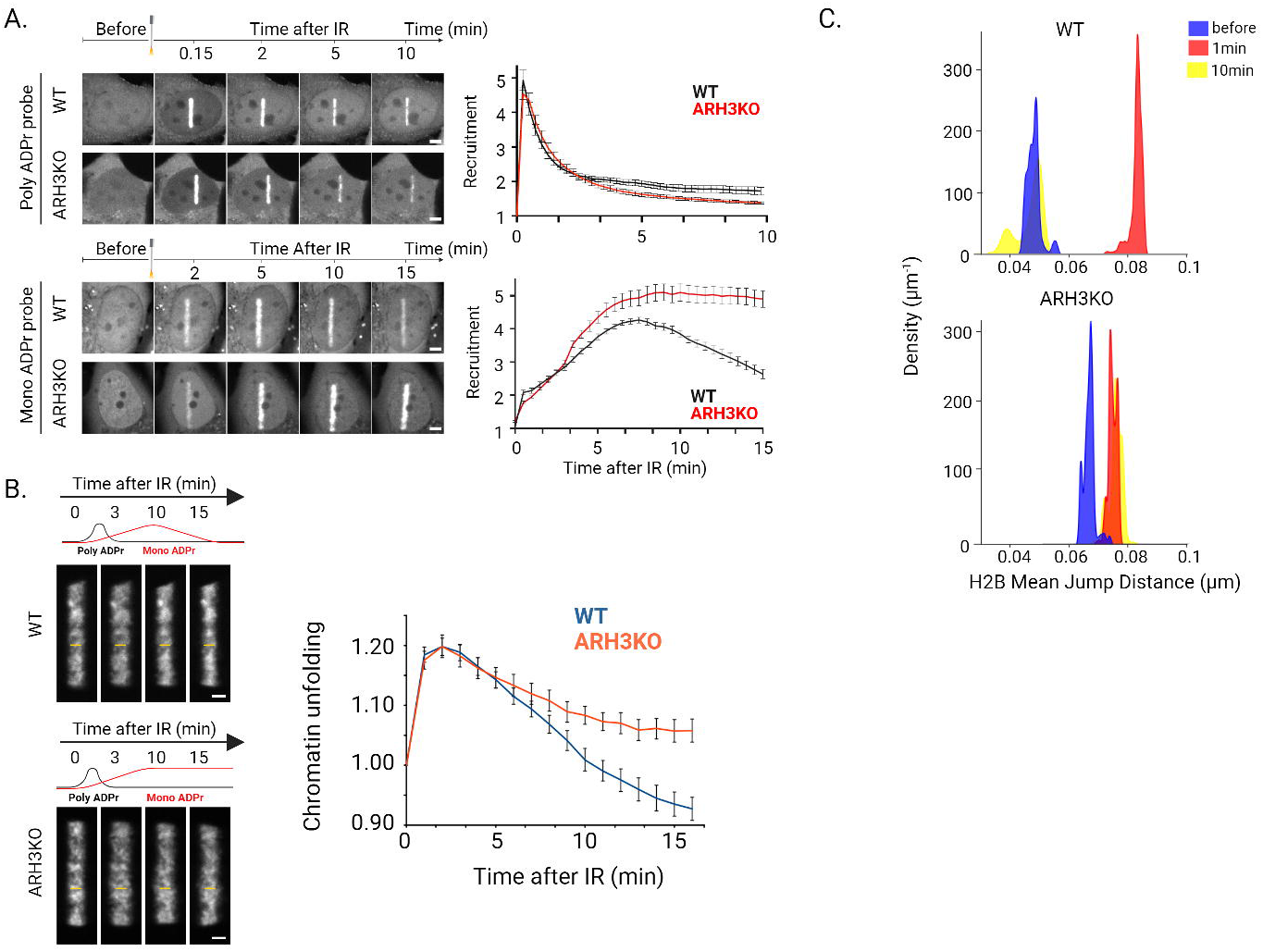
The removal of MAR marks by ARH3 is needed for the recovery of the chromatin state following its initial unfolding at sites of damage. (A) Representative confocal images and recruitment kinetics of the GFP tagged WWE domain of RNF146 (PAR sensor) and Macrodomain of Macro D2 (MAR sensor) expressed WT and ARH3 KO U2OS cells after irradiation at 405 nm. Scale bars: 4 µm. (PAR sensor N_WT_=12, N_KO_=12; MAR sensor N_WT_=11, N_KO_=12)(B) Representative confocal images and relative average thickness of the photo-activated damaged area in WT and ARH3KO cells expressing H2B-PAGFP and irradiated at 405 nm. Scale bars: 2 µm. Curves of the average thickness of the photo-activated line are mean ± SEM (WT N=12, ARH3 KO N=16). (C) Mean jump distance histograms for the immobile population of H2B tracks inside the irradiated region, before and at different times after micro-irradiation at 355 nm in WT and ARH3 KO U2OS cells. (WT N_pre_=12, N_1min_=4, N_10min_=3; ARH3 KO N_pre_=15, N_1min_=21, N_10min_=7).

## DISCUSSION

### Chromatin “breathing” at DNA lesions: a multiscale choreography of chromatin remodeling events

It is now well established that the DDR includes various chromatin remodeling steps which are crucial for the efficient and faithful restoration of genomic integrity^69^. In yeast, a compelling model has emerged in relation to double-strand break repair, where an increase in chromatin mobility triggered by H2A phosphorylation as well as the homologous recombination machinery facilitates homology search^29–32^. The picture remains less clear in mammals with different results depending on the time after damage induction as well as the chromatin landscape in which the lesions occur^3,23,28,70–73^. Importantly, most of these studies focused on a single spatial scale, which precludes the establishment of a global model for a chromatin structure that is inherently multiscale, sometimes even referred as fractal-like^74^. In this work, we aimed to overcome this technical limitation and enable comprehensive analyses of chromatin behavior by developing an original multiscale framework to assess early changes in the chromatin structure upon DNA damage at multiple levels: from the chromosome scale to the chromatin fiber and down to individual nucleosomes. We uncovered a “breathing” mechanism that affects the different folding scales of the chromatin immediately after damage induction (Figure 6). Our findings demonstrate the tight connection between chromatin mobility at the single nucleosome scale and its global compaction state, an aspect for which a unified general model was previously lacking, even beyond the DDR^44,56,75^. By monitoring the precise timing of this remodeling process, our work also reveals that the relationship between the different chromatin folding scales is more than a simple direct correlation. Indeed, while the rapid increase in the mobility of the nucleosomes along the fiber upon DNA damage is associated with a global unfolding, the maintenance of the resulting open chromatin state does not seem to require enhanced nucleosome dynamics. Therefore, the acute surge in nucleosome mobility appears as a transient “activated state” allowing for the switching between two persistent chromatin conformations displaying different compaction levels. Therefore, our work illuminates a sophisticated, multifaceted relationship between chromatin folding scales, paving the way for in-depth characterization of the mechanisms underlying the transitions between chromatin states.

**Figure 6.**
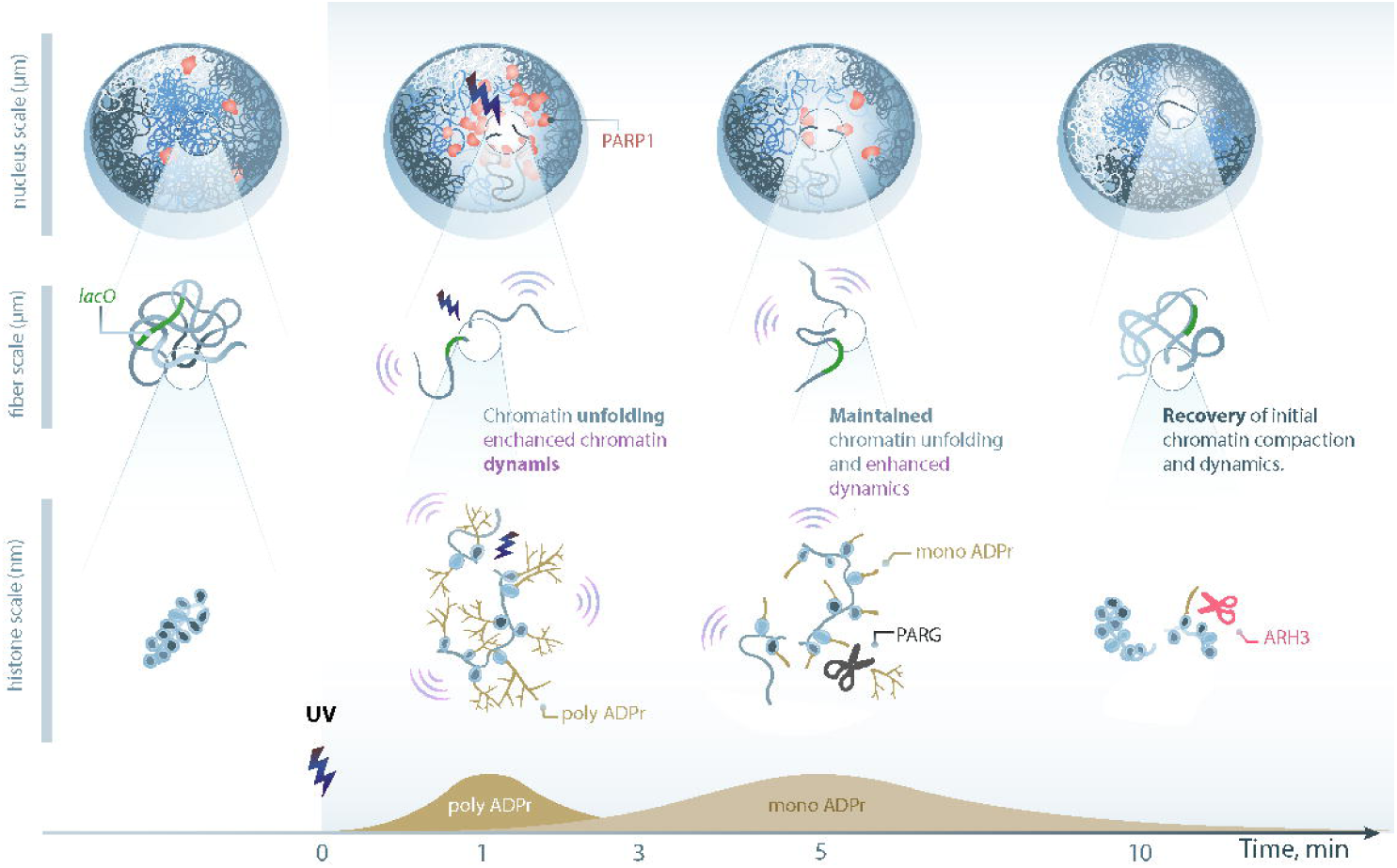
Model of ADPr-dependent multiscale chromatin breathing at sites of DNA damage. A few seconds after DNA damage, chromatin undergoes rapid unfolding along with an increase of its dynamics at the fiber to the nucleosome scale, a process triggered by histone ADP-ribosylation. While nucleosome dynamics rapidly drops, chromatin remains in an open state for several minutes until MAR erasing by ARH3 allows gradual recondensation. (Art by Olga Markova).

### Histone ADPr is both sufficient and necessary to promote multiscale chromatin unfolding at DNA lesions

Our current and previous findings^21^ demonstrate that decorating the chromatin fiber with ADP-ribose marks is itself sufficient to promote unfolding. This generic process, in coordination with the probably more specific activity of the multiple chromatin remodelers recruited to DNA lesions in an ADP-ribose dependent manner^4,67,76,77^, is crucial for the establishment of a repair-competent chromatin conformation in the vicinity of the DNA breaks. Our data, in line with *in vitro* results on isolated chromatin fibers, indicate that histone ADPr is unlikely to promote a major disruption of the nucleosome architecture leading to eviction and subsequent chromatin unfolding^4,19,78^. Rather, ADPr of linker histone was shown to inhibit its ability to promote chromatin compaction^79^. Therefore, ADPr may change the conformation of the nucleosome at the entry-exit site and lead to a partial eviction of linker histone^20^, thus promoting chromatin unfolding. Besides affecting nucleosome conformation, ADPr could also act at higher chromatin folding scales by inhibiting nucleosome self-association^78,80^. Negatively charged ADP-ribose chains on the nucleosomes may thus stiffen the chromatin fiber due to self-repulsion along the polymer, thus promoting reduced packing. This model is in line with the increased nucleosome mobility that we observed at sites of damage which, assuming that these motions can be described by a simple Rouse model, imply an increase in the rigidity of the chromatin fiber^32,48,81^. Given that inter-fiber nucleosome interactions were proposed to dominate over intra-fiber ones^82^ in the nucleus, histone ADPr may also impair fiber-fiber packing, leading to further decrease of the chromatin compaction state.

### A new role for MAR marks in maintaining chromatin in an open conformation at sites of DNA damage

The ADP-ribose signal at sites of DNA damage has historically been considered to be mainly composed of PAR polymers. However, recent technological advances have revealed prevalent MAR marks that exhibit kinetics different from those of PAR, implying distinct roles in PARP1 signaling ^9^. Our data indicate that, while the initial acute PAR wave might be necessary for the initial unfolding of chromatin, the more persistent MAR signal could be sufficient to maintain an open conformation in the vicinity of the DNA lesions (Figure 6). Therefore, our findings establish the first distinct functional role for this abundant and enduring signal generated by PARP1. The fact that the unfolding step is not associated with nucleosome disassembly implies that this reorganization can be easily reversed by the removal of the MAR signal along the chromatin fiber by MAR hydrolase ARH3. Since this recondensation phase leads to a chromatin compaction state that appears denser than before damage, the erasure of MAR marks may also be necessary for the addition of other modifications on the histone tails to promote a closed conformation. In favor of this hypothesis is the observation that ADPr competes with several other marks on histone tails^80,83,84^, with this competition potentially regulating certain aspects of the DDR^85^. Noteworthy, the open chromatin conformation induced by persistent MAR signal on histones may promote pathological unbalances in the transcriptional profiles of patient cells with ARH3 mutations associated with neurodegenerative diseases^86^.

### ADPr-dependent chromatin unfolding as a generic regulator of DNA accessibility

DNA wrapped around the nucleosomes shows higher susceptibility to MNase digestion upon histone ADPr^78^, suggesting that it became more accessible. At higher folding scales, nucleosome dynamics was proposed as a central regulator of chromatin accessibility in living cells^43,44^. Finally, we previously showed that the local relaxation of the chromatin controlled by ADPr at DNA lesions increases the binding rates of DNA-binding sensors^5^. Together, these findings draw a compelling picture in which the multiscale impact of histone ADPr on chromatin architecture triggers increased DNA accessibility in the vicinity of the lesions. While this might be a generic way to promote the accumulation of repair factors from different pathways at early stage of the DDR^21^, the subsequent recondensation regulated by ARH3 could potentially contribute to repair pathway choice due to the specific retention of a subset of these repair factors^39,67^. Besides the DDR context, our findings also demonstrate that a dynamic and accessible conformation may be a generic feature of ADP-ribosylated chromatin, independently of the presence of DNA lesions. Given that PARP1 also regulates transcription^87,88^, chromatin unfolding triggered by histone ADPr could facilitate access to transcription factors^88^. Therefore, our results identify the ADPr signaling as a key regulator of the dynamic chromatin conformation within the nucleus, potentially influencing multiple cellular functions involving DNA transactions.

## ACKNOWLEDGMENTS

The authors thank C. Hubert (Errol laser) for the installation of the micro-irradiation device on the single-molecule imaging setup. We thank C. Maison, D. Bailly, A. Forest (Institut Curie) and C. Fouquet (Institut de Biologie Paris Seine, Sorbonne University) for their help on the cell culture in the L2. The authors also thank the PICT-IBiSA@Pasteur Imaging Facility of the Institut Curie, a member of the France Bioimaging National Infrastructure (ANR-10-INBS-04), C. Chaumeton and F. Lam from the cellular Imaging Facility (Institut de Biologie Paris Seine, Sorbonne Université). We thank the Microscopy-Rennes Imaging Center (BIOSIT, Université de Rennes), member of the national infrastructure France-BioImaging supported by the French National Research Agency (ANR-10-INBS-04), for providing access to the imaging setups, as well as S. Dutertre and X. Pinson for technical assistance on the microscopes. We thank J. Morris, G. Timinszky, J. Ellenberg, S. Buratowski, A. Coulon and L. Lavis for sharing reagents. We are grateful to E. Fabre for her fruitful comments on the manuscript and A. Mansuy for sharing his software expertise. Illustrations in figures 2 and 4 created with BioRender.com. The S.H.’s and J.M.H.’s groups received financial support from the Agence Nationale de la Recherche (ANR-18-CE12-0015-03 RepairChrom and ANR-22-CE12-0039 AROSE). The J.M.H. team was financially supported by the programs iBio (Sorbonne University) and ATIP Avenir 2021. I.M.’s lab was funded by the Max Planck Society, the Deutsche Forschungsgemeinschaft (DFG, German Research Foundation) under Germanýs Excellence Strategy (CECAD, EXC 2030-390661388) and by the European Research Council (ERC-CoG-864117) to I.M. I.M. and E.J.L received support from the EMBO Young Investigator Program and the Cologne Graduate School of Ageing Research respectively. This work has inspired part of the Muse-IC project, a collaborative project between musicians and composers aiming to create musical pieces inspired by recent scientific discoveries.

## METHODS

### Plasmids

PmEGFP-PARP1, wild-type as well as the point mutants S499A/S507A/S519A (3SA) and LW/AA L1013A/W1014A (LW/AA), were previously described^21^, as well as pmEGFP-WWE and pmEGFP-Macrodomain of macroD221. pcDNA5/FRT/TO-FLAG-EGFP-BRCA1, pLacI-EGFP, p53BP1-EGFP and H2B-PAGFP were gifts from J.Morris^89^, G. Timinszky^90^ and J. Ellenberg^91^, respectively. To generate the pH2B-HaloTag, we amplified the HaloTag sequence from the plasmid pAT496 (pBS-SK-Halo-KanMX), kindly provided by C. Wu, using primers BshTI_ATG-Halo-Fwd (attaCACCGGTCGCCACCatggcagaaatcggtactgg) and NotI-Stop-End-Halo-Rev (attgcggccGCTTTAggaaatctctagcgtcgacagc) and replaced PAtagRFP in pH2B-PAtagRFP4 using BshTI / NotI.

### Cell culture

All cells used in this study were cultured in DMEM (Sigma) supplemented with 10% FBS, 100μgml^−1^ penicillin and 100Uml^−1^ streptomycin and maintained at 37°C in a 5% CO2 incubator. U2OS WT were obtained from ATCC. U2OS KO for PARP1 and ARH3 cells were kindly provided by I. Ahel^92^. The U2OS 2-6-3 cell line^93^ harbors a repetitive array of the *lacO* binding sequence at the chromosomal location 1p36 and was kindly provided by A.Coulon. For transient expression, the GFP-tagged plasmids were transfected with X-tremeGENE HP (Sigma) according to manufacturer instructions. To establish cell lines stably expressing H2B-HaloTag, cells were transfected with the H2B-HaloTag plasmid and selected using media supplemented with 500 μg.ml^−1^ G418. The PARP1 inhibitors Talazoparib (Euromedex) were used at 30 µM and added to the cell medium 10 minutes prior imaging. For PARG inhibition, cells were treated with 25 µM of PDD00017273 (Bio-Techne, USA) for the indicated durations. For HaloTag labeling, cells were incubated for 30 minutes with 10 nM of HaloTag ligands conjugated to the photoactivatable dye JF549, kindly provided by L. Lavis. For Hoechst presensitization, cells were bathed with culture medium containing 0.3 μg/ml Hoechst 33342 (Sigma) for 1 h. Immediately before imaging, growth medium was replaced with CO_2_-independent imaging medium (phenol red-free Leibovitz’s L-15 medium, ThermoFisher). All live-cell experiments were performed on unsynchronized cells.

### Western blotting

Cells were lysed on Triton-X buffer (1% Triton X-100, 100 mM NaCl, 50 mM Tris-HCl, pH 8.0, 5 mM MgCl2, 0.1% Benzonase (Sigma-Aldrich), 1× protease inhibitor (Roche)) on an orbital rotator for 30 min at 4 °C. Samples were centrifuged at 20,000g for 15 min, and supernatant was collected. Protein samples were quantified using Bradford (Bio-Rad), and equal amounts of protein were loaded on gels for SDS–PAGE prior to immunoblotting. The membranes were blocked in PBS buffer with 0.1% Tween20 and 5% non-fat dried milk for 1h at room temperature and incubated overnight at 4°C with the following primary antibodies: anti-pan-ADPr (MABE1016, Sigma, 1:1500), which binds both MAR and PAR marks, anti-PARP1 (homemade^4^, 1:10000), anti-ARH3 (hpa027104, Sigma, 1:1500), anti-H3 (Ab1731, Abcam, 1:2500). Then, the membranes were incubated with a peroxidase-conjugated secondary anti-rabbit antibody (P039901-2, Agilent, 1:3000) for 1h. Blots were developed using ECL (Thermo) and analyzed by exposing to films.

### Confocal imaging and quantification

Changes in the chromatin compaction state and protein recruitment at sites of laser irradiation was performed as previously described^4,21^. In brief, images were acquired either on a Ti-E inverted microscope from Nikon equipped with a CSU-X1 spinning-disk head from Yokogawa, a Plan APO 60x/1.4 N.A. oil-immersion objective lens and a sCMOS ORCA Flash 4.0 camera for Hamamatsu; or on an Olympus Spin SR spinning disc system equipped with a CSU-W1 spinning-disk head from Yokogawa (50 micron pinhole size), a UPLSAPO 100XS/1.35 N.A. silicon-immersion objective lens and a sCMOS ORCA Flash 4.0 camera. Laser irradiation of Hoechst-presensitized cells was performed along a 10 or 16 µm-line through the nucleus with a continuous 405 nm laser set at 125-130 mW at the sample level. Recruitment of GFP tagged BRCA1 was monitored on a Zeiss LSM 880 confocal setup equipped with a C-Apo ×40/1.2 N.A. water-immersion objective and a GaAsP detector array for fluorescence detection. The pixel size was set to 80 nm. Nuclei of non-sensitized cells were irradiated within a region of interest of 100-pixel width and 10-pixel height with a Ti:sapphire femtosecond infrared laser (Mai Tai HP, Spectra-Physics) with emission wavelength set to 800 nm. For all these live-cell imaging experiments, cells were maintained at 37°C with a heating chamber. The changes in the chromatin compaction state were measured using a custom MATLAB routine that estimates the thickness of the photo-converted H2B line relative to its value immediately after damage induction. To quantify protein recruitment, the mean fluorescence intensities were evaluated over time within the irradiated area and the whole nucleus, both segmented manually on ImageJ/FIJI or Olympus CellSense. After background subtraction, the intensity in the irradiated area was divided to the nuclear intensity to correct for imaging photobleaching, and then normalized to the signal prior to DNA damage.

### Single-particle tracking

Dynamics of the LacO array and the single H2B proteins were monitored on an inverted Nikon Ti microscope, equipped with an EM-CCD camera (Ixon Ultra 897 Andor) and a 100x/1.4NA or 1.3 NA oil-immersion objective, leading to a pixel size of 160 nm. Cell were maintained at 37°C using a Tokai device (STXG-TIZWX-SET). A pulsed diode 355 nm laser monomode remotely controlled with the Pangolin Software (LASER ERROL) was coupled to the microscope to allow laser irradiation within a predefined line within the cell nucleus (3.2µm x 0.4µm) for 110 ms. The dynamics of the *lacO* array was monitored at a frame rate of 33 Hz. Fluorescent beads (FluoSpheres, ThermoFisher) were used as fiducial markers to correct for cell drift. For the tracking of single H2B-Halo in living cells, the PA-JF549 ligand was photo-activated by the 405 nm laser (1 pulsation every 10 frames, power of 0.006 kW/cm^2^ at the sample) and excited by the 561 nm laser (continuous excitation, power of 7 kW/cm^2^ at the sample). Single molecule imaging sequences of 5000 frames were acquired at a frame rate of 100 Hz. For single molecule detection, position refinement and track reconstruction, we used the SlimFast multi-target tracking algorithm^94^ with the following parameters: localization error: 10^-6^; deflation loops: 0; max OFF time: 1; max D: 7 μm^2^/s. Home-made routines written in Matlab (Mathworks) were used to visualize the detection density maps and trajectories. H2B dynamics was measured within a rectangle of 3 μm large and whose height was limited by the nucleus border, which was either encompassing the irradiated area or localized away from it. Approximately 1 000 trajectories per imaging sequence were monitored within such region of interest, allowing for the building of the mean jump distance histograms. The mean single-molecule track length was approximately 150 ms (15 frames), much shorter than the characteristic fluorescence decay of 12.5 s estimated for the PA-JF549 ligand. Therefore, track lengths are limited by out-of-focus movement rather than PA-JF549 photobleaching.

### Mean squared displacement analysis

To characterize the dynamics of the *lacO* array, the time-average mean squared displacement curves were derived from each trajectory as follows:

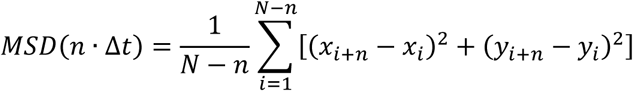

where N is the total number of points within the trajectory, (x, y) the coordinates of the locus in 2-dimensions and Δt the time interval used during the acquisition. To obtain a precise estimation of the fitted parameters, we calculated time-ensemble-averaged MSD over several trajectories, which are simply referred to as “MSD” in the Figures using a home-made Matlab code^48^. In line with previous studies^48,81,95^, the mean MSD curves were fitted with the following anomalous diffusion model:

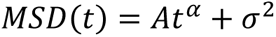

where *α is* the anomalous exponent, *A* the anomalous diffusion coefficient and *σ* the positioning accuracy. Here, we found a better agreement with the anomalous diffusion model, consistent with previous studies ^48,81,95^. *α* <1 correspond to subdiffusive dynamics, referring to tracked objects that reiteratively scans neighboring regions before reaching a distant position^96^. In contrast, *α* >1 correspond to motions displaying a directed component^97^. The anomalous diffusion coefficient *A* quantifies motion amplitude. It is proportional to the diffusion coefficient only in the case of normal diffusion (*α* = 1), which is rarely observed in biological systems. Besides this analysis of diffusion anomality, we also quantified locus mobility with an effective diffusion coefficient D*_lacO_* calculated as D=p/4 where p is the slope of the linear fit of the first 4 points of the MSD curves. To compare with the diffusion of the slow population of H2B, we also extracted an effective diffusion coefficient D_H2B_. Since H2B trajectories are much shorter than *lacO*, to reduce the experimental and localization noise, we calculated a denoised averaged diffusion coefficient by measuring the tangents of the fitted MSD curve between 0.1s and 1.0s.

### Classification of the H2B tracks

We used a convoluted neural network (CNN)^98,99^ to classify H2B trajectories in 3 categories: immobile, hybrid and mobile (Figure S1E-G). In the preprocessing step, the single-molecule trajectories are converted into 2D images of 512×512 pixels to consider the sub-pixel accuracy of localization. Displacements between two consecutive positions are interpolated as a straight segment. Each segment is given a third dimension, defined as a color, according of the instantaneous diffusion coefficient calculated between the corresponding two positions: 0 μm^2^/s < red ≤ 0.5 μm^2^/s, 0.5 μm^2^/s < green ≤ 1 μm^2^/s, 1 μm^2^/s < blue respectively. Starting from 1040 images of tracks, data augmentation by image rotation was used to generate a set of 71,760 images. After shuffling, 80% of these tracks were composing our training set while the remaining 20% were used as a validation set. The model is composed of 5 CNN layers after max-pooling, with a last dense layer with three outputs and softmax activation. The model is trained with the Adam optimizer based on the cross-entropy loss. The trained model shows around 98% accuracy on the validation set. Data is shown in mean jump distance histograms that indicate the distance a molecule travels in a given space and time interval, as a function of the probability density per unit length.

## Notes

### Competing Interest Statement

The authors have declared no competing interest.

