## Supplementary information for "Single-nucleosome imaging reveals principles of transient multiscale chromatin unfolding triggered by histone ADP-ribosylation at DNA lesions"

**Garcia-Fernandez *et al*.**

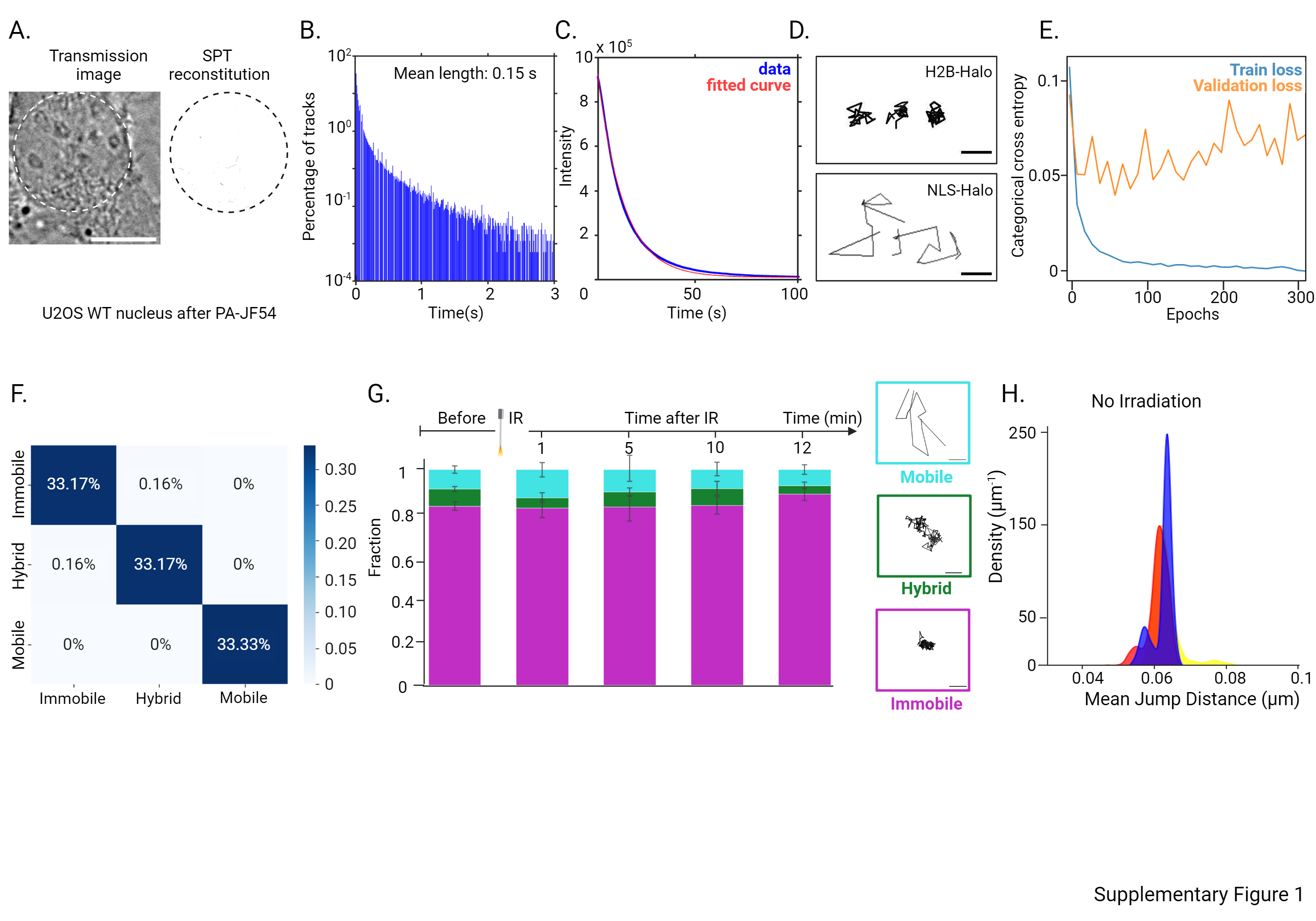

**Figure S1. Tracking of single fluorescently tagged H2B proteins in living cells.** (A) Control U2OS cell that does not express H2B-Halo after tagging with PA-JF549 Halo ligand. The widefield transmission image of the nucleus (highlighted by the dotted line) is shown on the left and the single molecule trajectories that were detected is shown on the right. (B) Distribution of tracks length for U2OS cells expressing H2B-Halo and tagged with PA-JF549 Halo ligand in the absence of DNA damage. The histogram combines 60 cells representing 68779 trajectories (mean length of 0.15s). (C) Characteristic bleaching curves for PA-JF549 Halo ligand tagging H2B-Halo expressed in U2OS cells. The experimental data (blue) were fitted with the single exponential decay (red) to recover a characteristic bleaching time of 12.3 s (n=5). (D) Representative single molecule trajectories for H2B-Halo (top) and NLS-Halo (bottom) tagged with PA-JF549 Halo ligand. (E) Training and validation cross entropy losses for the CNN model used to classify the single molecule H2B trajectories. Minimum for the validation set is reached at 6 epochs. (F) Confusion matrix of the trained model on the validation set of H2B trajectories. (G) Distribution of the H2B trajectories obtained from the CNN classifier at different time points after irradiation at 355 nm. Magenta, green and cyan bars represent immobile, hybrid and mobile H2B populations, respectively. Representative example of trajectories are shown on the right. Scale bars: 100nm. (G) Mean jump distance histograms for the immobile population of H2B tracks in the absence of micro-irradiation (N_bef_=4, N_1min_=4, N_10min_=4).

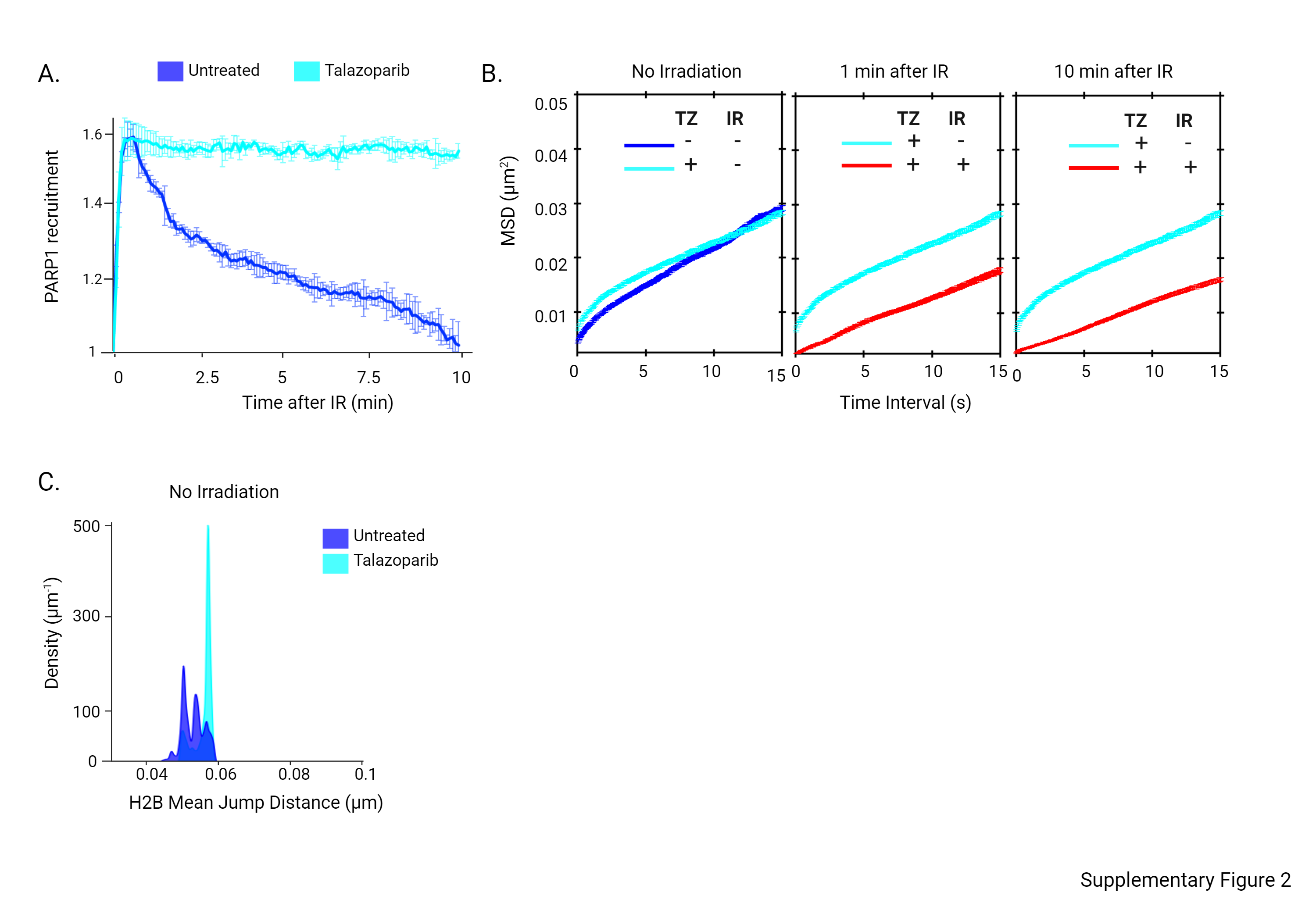

**Figure S2. Impact of PARP inhibition on chromatin dynamics.** (A) Kinetics of PARP1 recruitment to DNA damage induced by 355 nm irradiation in WT U2OS cells left untreated or treated with 30 µM of Talazoparib (N=10 for each condition). (B) Mean squared displacement curves of the fluorescently tagged *lacO* array in cells treated or not with 30 µM Talazoparib, before, 1 min and 10 min after irradiation (N=10). (C) H2B mean jump distance histograms for the immobile population of H2B tracks in WT cells treated or not with 30 µM Talazoparib in the absence of DNA damage (untreated WT N=75, WT+Talazoparib N=52). Mean jump distance between Talazorib-treated condition *versus* untreated condition is significantly different (p < 0.001, calculated from Yuen-Welch Test).

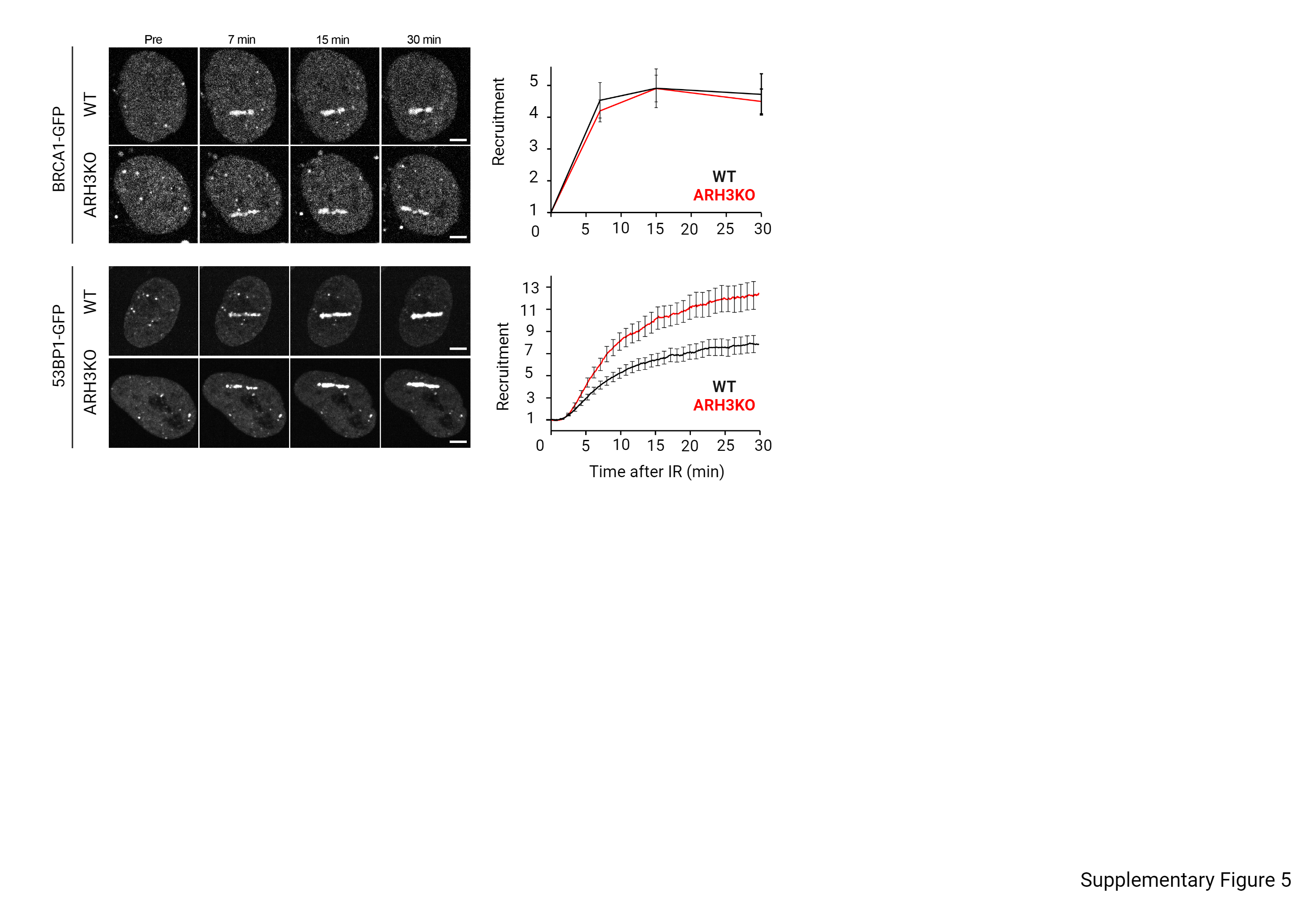

**Figure S3. Impact of the loss of ARH3 on the recruitment of repair factors.** Confocal image sequence and recruitment kinetics of BRCA1-GFP and 53BP1-GFP in WT and ARH3 KO U2OS cells after irradiation at 800 nm or 405 nm, respectively. Scale bars: 4 µm. Data are shown as mean ± SEM. (BRCA N_WT_=30, N_ARH3_=32; 53BP1 N_WT_=18, N_ARH3_=18).
